# Comparison of different orders of Legendre polynomials in random regression model for estimation of genetic parameters and breeding values of milk yield in the Chinese Holstein population

**DOI:** 10.1101/562991

**Authors:** Jianbin Li, Hongding Gao, Per Madsen, Wenhao Liu, Peng Bao, Guanghui Xue, Jun Yang, Yundong Gao, Guosheng Su

## Abstract

Random regression test-day model has become the most commonly adopted model for routine genetic evaluations for different dairy populations, which allows accurately accounting for genetic and environmental effects at different periods during lactation. The objective of this study was to explore appropriate random regression test-day model for genetic evaluation of milk yield in Chinese Holstein population. Data included 419,567 test-day records from 54,417 cows in the first lactation. Variance components and breeding values were estimated using random regression test-day model with different order (first order to fifth order) of Legendre polynomials, and accounted for homogeneous or heterogeneous residual variance across the lactation. The goodness of fit of the models was evaluated by total residual variance (TRV) and − 2*logL*. Further, the predictive ability of the models was assessed by Spearman’s rank correlation between estimated breeding values for 305d milk yield (EBV_305_) from the full data set and reduced data set in which the records from the last calving year were masked. The results showed that random regression models using third order Legendre polynomials (LP3) with heterogeneous residual variance achieved the lower TRV and − 2*logL* value and the highest correlation for EBV_305_ between full data and reduced data. Heritability estimated by this model was 0.250 for 305d milk yield and ranged from 0.163 to 0.304 for test-day milk yield. We suggest random regression model with Legendre polynomial of order 3 and accounting for heterogeneous residual variances could be an appropriate model to be used for genetic evaluation of milk yield for Chinese Holstein population.

## INTRODUCTION

In China, the dairy herd improvement project (DHI) was firstly implemented in 4 provinces in the 1990s. In 2006, the Ministry of Agriculture of China approved a project to promote DHI project in 8 provinces where there were many large dairy populations (1). Lately, the project has been expanded to 25 provinces in China, where provincial DHI laboratories and data centers have been established (2), and there were about 700,000 cows recorded milk production in China each year.

Random regression test-day model has been widely used in genetic evaluation for production traits in dairy cattle, which has many advantages including more accurately accounting for genetic and environment effects at different stages of the lactation, thus resulting in more reliable genetic evaluation (3–7). It has been reported that test day model is significantly better than lactation model (using full and extended 305d lactation records) with 2-3% increase in accuracy for bulls and 6-8% for cows for milk yield first lactation (8). In addition, test-day model allows to predict estimated breeding value (EBV) for each test day, each particular period or the full lactation (305d) (7).

The functions generally used to model the lactation curve include Woods’s model (9), Wilmink’s function (10), Spline function (Pelmus,R.S.,et al. 2016) and Legendre polynomial function (11). Because of differences in production environments and management systems, optimal functions for test models in different countries may be different (12, 13). Several studies have shown that Legendre polynomials (LP) fit random regression test-day model well in general, but there is no “gold standard” reported in literatures on choosing optimal order of LP in the model, and the choice of the order of fit is highly depending on the practical data structures. For example, a fourth order LP (LP4) was used for national genetic evaluations in Canada and Italy, and a fifth order LP (LP5) was used in UK (14). The joint Nordic test-day model is a multivariate model for milk, protein and fat in lactation 1 to 3 (in total 9 traits). The genetic and permanent environment effects are modeled by second order LP extended with an exponential term *e*^−0.04 × *DIM*^(15).

Although a large number of new milk records are collected monthly in China, routine genetic evaluation has not been performed timely. In addition, genetic parameters are not updated regularly. To use data efficiently and reduce the cost of keeping candidate bulls, it is necessary to perform genetic evaluation frequently, such as, genetic evaluation is performed five times per year by Interbull (16). Moreover, there has been no study to investigate the impact of parametric functions for lactation curve on genetic evaluation in Chinese Holstein population.

The aim of this study was to find an optimal order of Legendre polynomials for genetic evaluation of milk yield in Chinese Holstein population by comparing different orders of Legendre polynomials in random regression test day models in terms of goodness of fit and prediction accuracy.

## MATERIALS AND METHODS

### Data

Data were obtained from the database at Dairy Cattle Research Centre DHI Lab, Shandong Academy of Agricultural Sciences. First lactation records from 2004-2015 which fulfill the following criteria: ages at calving between 20 and 38 months, and daily milk yield between 5 and 80kg, days in milk (DIM) between 5 and 305, and cows with at least 3test day records were extracted. The final data consisted of 419,567 test day records from 54,417 cows. Number of records in each DIM class ranged from 576 to 1,768. Descriptive statistics of the data are presented in Table 1. The traced pedigree included 104,884 individuals.

**Table 1.**
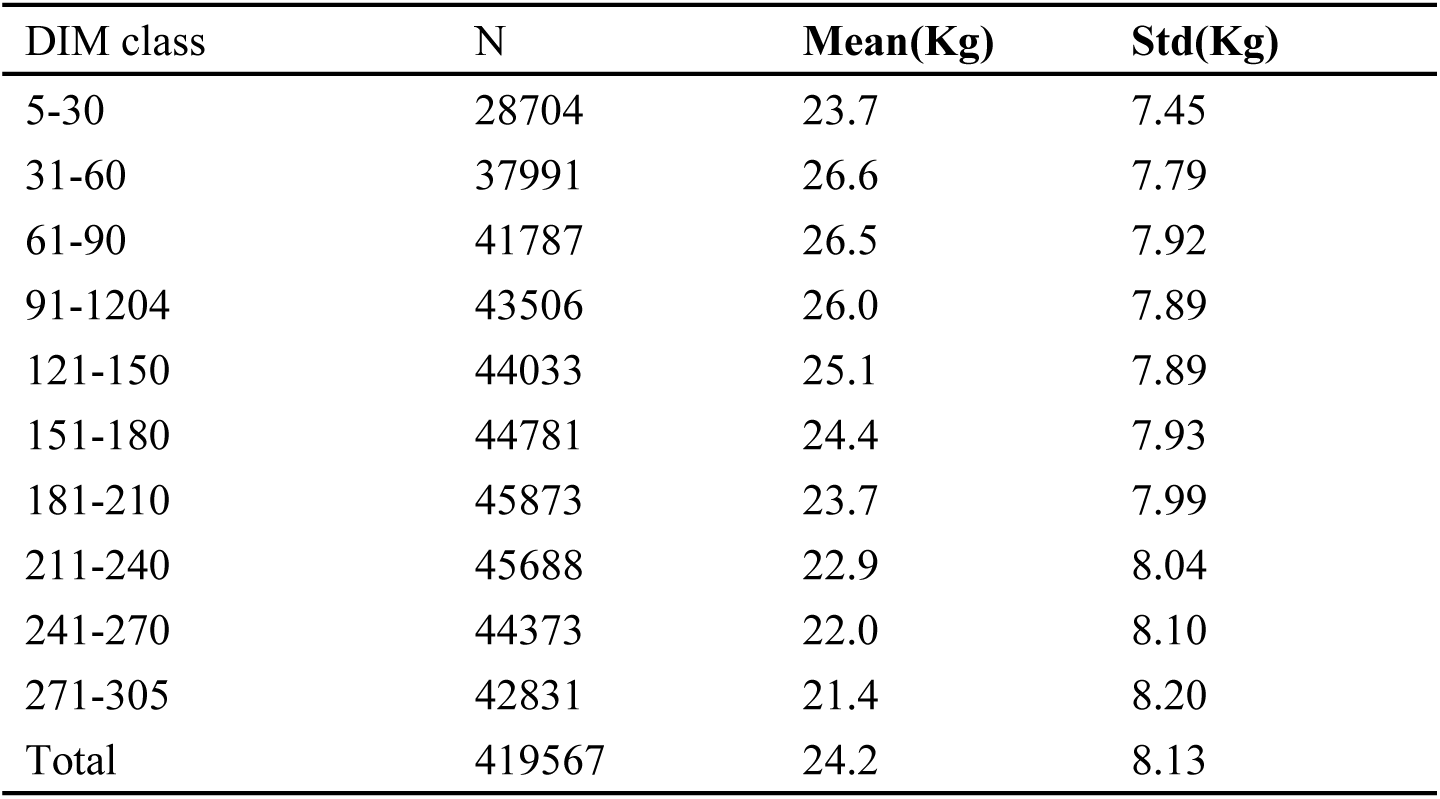
Number of records in each days in milk (DIM) class (N), Mean, Standard deviation (Std)

| DIM class | N | Mean(Kg) | Std(Kg) |
| --- | --- | --- | --- |
| 5-30 | 28704 | 23.7 | 7.45 |
| 31-60 | 37991 | 26.6 | 7.79 |
| 61-90 | 41787 | 26.5 | 7.92 |
| 91-1204 | 43506 | 26.0 | 7.89 |
| 121-150 | 44033 | 25.1 | 7.89 |
| 151-180 | 44781 | 24.4 | 7.93 |
| 181-210 | 45873 | 23.7 | 7.99 |
| 211-240 | 45688 | 22.9 | 8.04 |
| 241-270 | 44373 | 22.0 | 8.10 |
| 271-305 | 42831 | 21.4 | 8.20 |
| Total | 419567 | 24.2 | 8.13 |

**Table 2.**
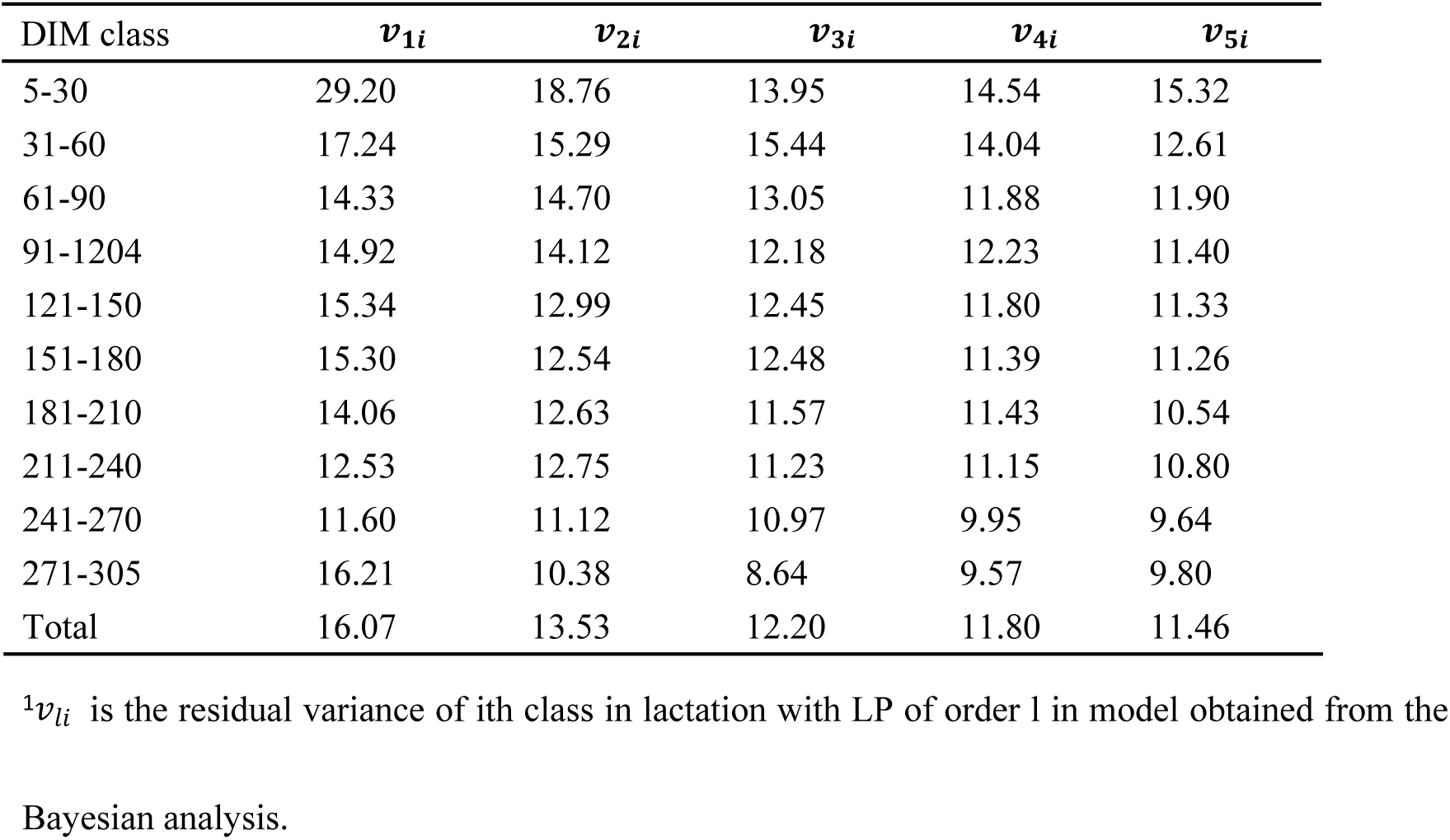
Estimated residual variances (*v*_*li*_)^1^ for different classes of days in milk and different Legendre polynomials

### Model

We used first order to fifth order Legendre polynomial (LP1 to LP5) to fit random regression test day model, respectively. The model equation was as follows:

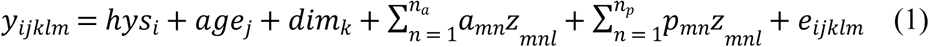

Where *y*_*ijklm*_ is the observation within ith herd-year-season effect, the jth age classes, the kth DIM effect on lth test day of cow *m*; *hys*_*i*_ is the ith fixed herd-year-season effect; *age*_*j*_ is the jth fixed calving age effect; *dim*_*k*_ is the kth fixed DIM effect; *a*_*mn*_ is the nth random regression coefficients for additive genetic effect of cow m; *p*_*mn*_ is the nth random regression coefficients for permanent environmental effect of cow m; *z*_*mnl*_ is Legendre polynomials on DIM, and *n*_*a*_, *n*_*p*_ are orders (from 1 to 5) of Legendre polynomials for additive genetic and permanent environmental effects, respectively; and *e*_*ijklm*_ is residual effect.

In this study, March, April, May, September, and October were defined as calving season1, June, July, and August as calving season2, November, December, January, and February as calving season3, and there were 1,891 herd-year-season classes in total. Calving age was classified into 4 levels: 20-23mo, 24-27mo, 28-31mo, 32mo or later. Residual variance was assumed either homogeneous or heterogeneous across lactation. For models with heterogeneous residual variances, residuals were divided into10 classes (5-30, 31-60, 61-90, 91-120, 121-150, 151-180, 181-210, 211-240, 241-270, and 270-305 DIM) (17). Bayesian method with Gibbs sampling was used to generate the posterior samples for models with heterogeneous residual variances. The length of MCMC chain was set to 55 000 with a burn-in of 5 000 iterations. Convergence diagnostics for MCMC were assessed using R package boa (18) and all parameters investigated had converged to the posterior distribution. The estimates of residual variances for different periods are shown in Table2. To compare models with different orders of Legendre Polynomials, the heterogeneous residual variances were handled by putting different weights on residual variance for different periods of DIM. The weights were calculated by 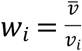, where *v*_*i*_ is the posterior means for residual variance of ith DIM class obtained from the Bayesian analysis, 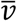 is the mean of residual variances.

Additive genetic variance for a particular DIM was calculated as 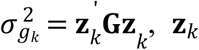 is a column vector of LP coefficients at kth DIM, **G** is covariance matrix of additive genetic effect. Permanent environmental variance for a particular DIM was 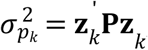, matrix **P** is covariance matrix of permanent environmental effect, **z**_*k*_ is same as above; EBV of a particular animal at a particular DIM was calculated as 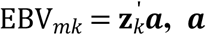, **a** is column vector of additive genetic random regression coefficients of a particular animal, **z**_*k*_ is same as above; The EBV for the whole lactation was calculated as 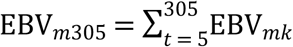. The estimation of variance components and prediction of breeding values using different models were carried out by the DMU package (19).

### Model comparison

Models with different orders of Legendre polynomials were compared using following methods based on full and reduced data sets:

a. Residual variance of 305d milk yield 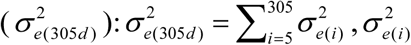 is residual variance of each TD, which is the same in each TD when considering homogeneous variance, however, different but same in each class in lactation when considering heterogeneous variance. A smaller 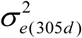 indicates a better fitting of the regression model.
b. Log-likelihood ratio test (LRT) was used to test the differences between the reduced order model and the subsequently augmented model with addition of one extra order (LP1 vs. LP2, LP2 vs. LP3, LP3 vs. LP4, LP4 vs. LP5). The LRT between models with successive order is *LRT*_*i*_ = − 2*logL*_*i*_ − (− 2*logL*_i+1_)with *df*_*i*_ = *nP*_i+1_ − *nP*_*i*_. In this study, − 2*logL*_i_ is − 2*logL* value for the model with ith order, *nP* is the number of parameters in the corresponding model.
c. Spearman’s rank correlation: It was used to evaluate predictability of model that the spearman’s rank correlation between EBV_305_ of animals calving in last year in data, whose EBVs were estimated based on including their own phenotypes and masking their phenotypes. The correlation was calculated as 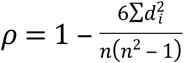 (Bolboac, S.D.& Lorentz J. 2006), *d*_*i*_ is the difference between the two ranks, *n* is the number of cows.

## RESULTS

### General statistics of TD milk

Mean for TD milk yield in the different class of lactation was showed in Table 1, where means ranged from 21.4 kg to 26.6 kg with standard deviations from 7.45 kg to 8.20 kg. An increase in milk yield was found up to 53 DIM, followed by a gradual decrease until the end of lactation. Averaged over different classes, TD milk was 24.2 kg with a standard deviation of 8.13 kg in first lactation Holstein cows.

### Goodness of fit

Table 3 presents the estimated parameters using random regression test-day model based on different assumption of residual variances (homogeneous or heterogeneous). Number of parameters for random effects of models was increased from 7 to 43 when increasing the order of LP from LP1 to LP5. The estimated residual variances 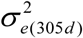 were decreased from 4661.46 to 3354.87 as order increased when assuming homogeneous residual variance, and from 4557.65 to 3273.39 when assuming heterogeneous residual variances. The differences between LP1 and LP5 were 1306.59 for homogeneous residual variance and 1284.26 for heterogeneous residual variance. However, the differences were smaller between LP3 and LP4 and between LP4 and LP5, which were 171.43 and 142.08 for homogeneous, 132.99 and 169.56 for heterogeneous variances, respectively. Differences of 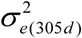 between models with homogeneous and with heterogeneous residual variances were 103.81, 60.03, 92.44, 54.00 and 81.48 corresponding LPs (from LP1 to LP5) respectively. That means 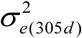 were decreased when considering heterogeneous residual variances.

**Table 3.** Number of additive genetic effect (nA) and permanent environment effect (nPE), and estimates of residual variance of cumulative 305d milk yield 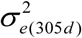, *-2*LogL, the log-likelihood ratio test (LRT) between the reduced order model and the subsequently augmented model with addition of one extra order, degree of freedom (df) and *χ*^2^ of LRT

| | LP | nA | nPE | $\sigma_{e(305d)}^2$ | -2LogL | LTR<sup>3</sup> | df | $\chi_{0.005}^2$ |
| --- | --- | --- | --- | --- | --- | --- | --- | --- |
| HO<sup>1</sup> | LP1 | 3 | 3 | 4661.46 | 1757621.90 | 18462.74* | 6 | 18.548 |
|  | LP2 | 6 | 6 | 3985.43 | 1739159.16 | 7013.04* | 8 | 21.955 |
|  | LP3 | 10 | 10 | 3668.38 | 1732146.12 | 3292.53* | 10 | 25.188 |
|  | LP4 | 15 | 15 | 3496.95 | 1728853.59 | 2057.64* | 12 | 28.299 |
|  | LP5 | 21 | 21 | 3354.87 | 1726795.95 | - | - | - |
| HE<sup>2</sup> | LP1 | 3 | 3 | 4557.65 | 1725335.19 | 3325.47* | 6 | 18.548 |
|  | LP2 | 6 | 6 | 3925.40 | 1722009.72 | 1387.18* | 8 | 21.955 |
|  | LP3 | 10 | 10 | 3575.94 | 1720622.54 | 4376.24* | 10 | 25.188 |
|  | LP4 | 15 | 15 | 3442.95 | 1716246.30 | 2702.95* | 12 | 28.299 |
|  | LP5 | 21 | 21 | 3273.39 | 1713543.35 | - | - | - |

<sup>1</sup> HO was homogeneous residual variance; <sup>2</sup>HE was heterogeneous residual variance,
<sup>3</sup>LRT between LP1 and LP2, LP2 and LP3, LP3 and LP4, LP4 and LP5
\* P<0.005

*-2LogL* were decreased from 1757621.90 to 1726795.95 as order increased when assuming homogeneous residual variance, *-2LogL* decreased from 1725335.19 to 1713543.35 when assuming heterogeneous residual variance. The differences between models tested by Chi-square statistic of LRT were significant (P<0.005). Thus, the null hypothesis of equality of models with different orders was rejected. Differences of *-2LogL* between models with homogeneous and with heterogeneous residual variances were 32286.71, 17149.44, 11523.58, 12607.29 and 13252.6 corresponding LPs (from LP1 to LP5). That means *-2LogL* were decreased when considered heterogeneous residual variances.

### Comparison on estimated variances

Figure 1 and Figure 2 show the genetic variances 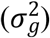, permanent environmental variances 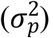, residual variances 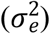, heritabilities (*h*^2^) and repeatabilities (*Rep*) at each TD along the lactation calculating based on the estimated covariance function coefficients. The curve of 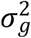 for TD showed a sharp decreasing in early lactation and then increasing from middle to the end of lactation.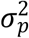 for TD presented similar trends as 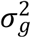. Estimates of 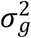 and 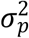 were somewhat different in models with different orders. 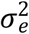 for TD was same when considering homogenous in same model, and not continuous when considering heterogeneous, however, they decreased with the order increasing. For heterogeneous variance, residual variance decreased from the beginning to the end lactation stage. The curve of heritability and repeatability showed a sharp decrease in early lactation and then increased from middle to the end of lactation.

**Figure 1.**
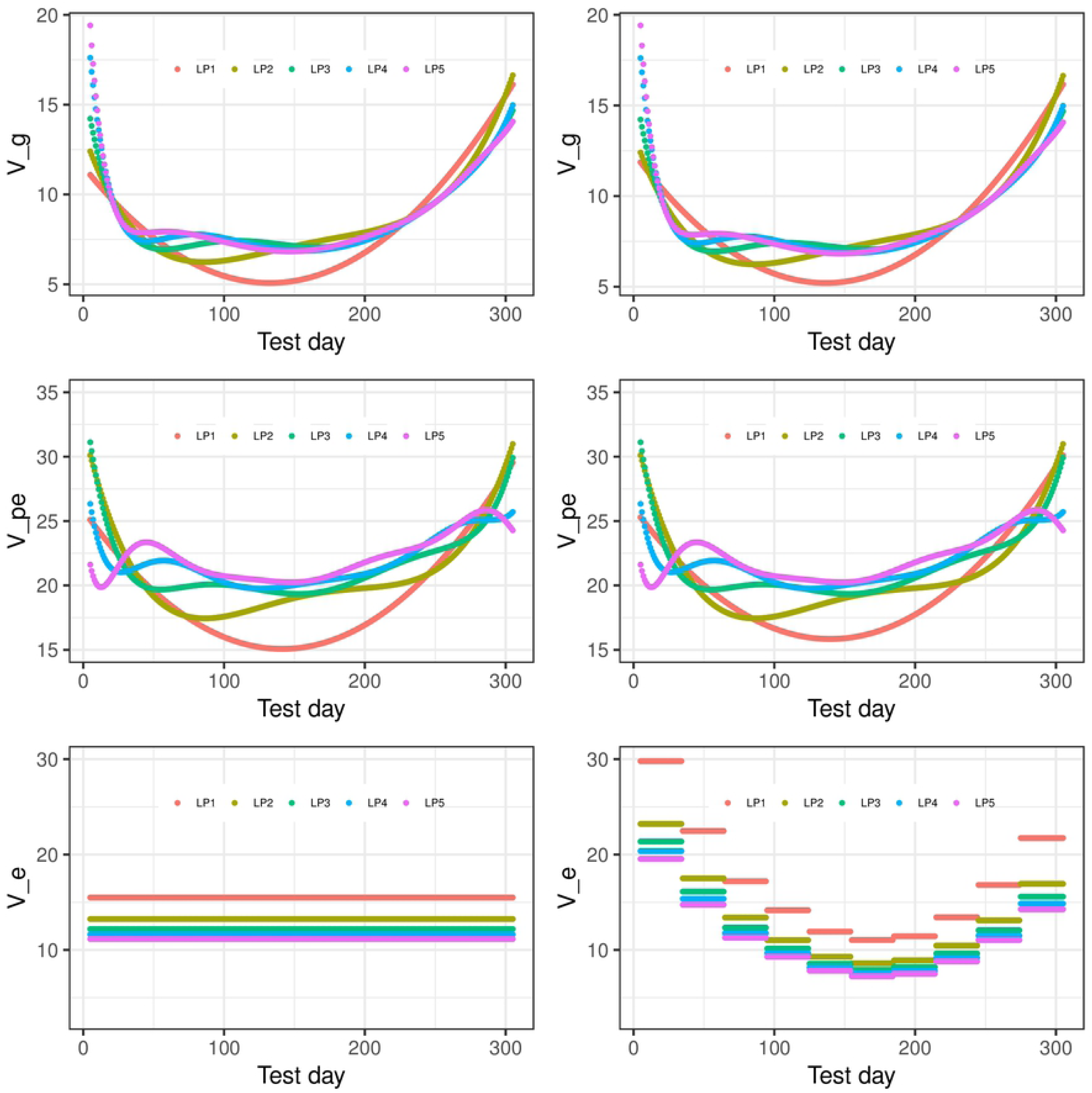
Genetic variances (V_g), permanent environmental variances (V_pe), residual variances (V_e) at each test day along the lactation from models with different orders of Legendre polynomials (LP) based on assumption of homogeneous residual variance (left column) or assumption of heterogeneous residual variance (right column)

**Figure 2.**
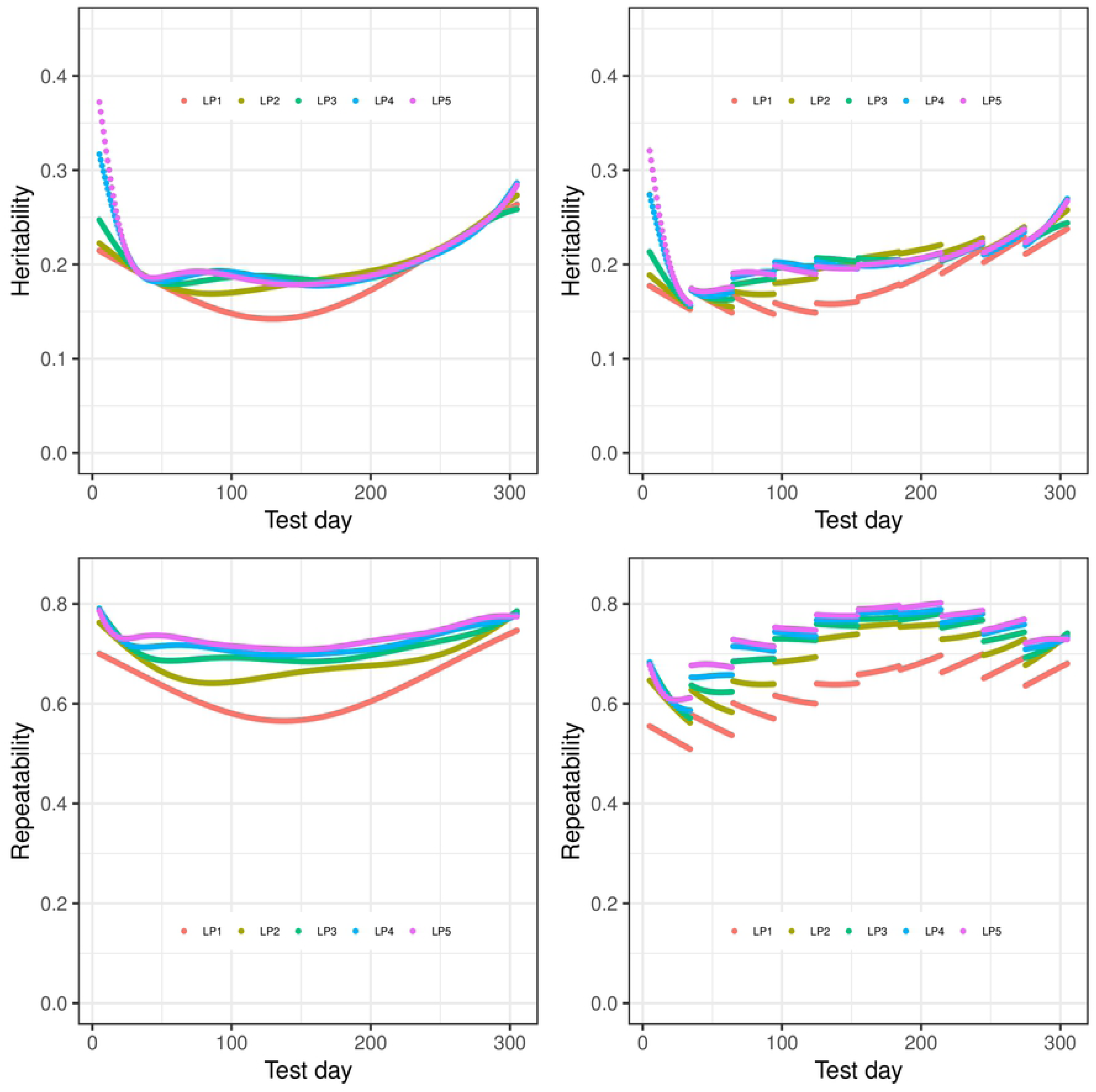
Heritabilities and repeatabilities at each test day along the lactation from models with different orders of Legendre polynomials (LP) based on assumption of homogeneous residual variance (left column) or assumption of heterogeneous residual variance (right column)

Table 4 shows the heritability (*h*^2^) and repeatability (*Rep*) for 305d milk yield and minimum and maximum values of TD milk yield. *h*^2^ for 305d estimated from models with different LPs ranged from 0.250 to 0.257 and *Rep* were from 0.741 to 0.749 when considering homogeneous variance. When considering heterogeneous variance, *h*^2^ for 305d were between 0.250 to 0.260, and *Rep* were between 0.738 and 0.749. *h*^2^ for TD from models with different LPs ranged from 0.142 to 0.372 and *Rep* were between 0.566 and 0.786 when considering homogeneous variance. When considering heterogeneous residual variances, *h*^2^ for TD ranged from 0.143 to 0.326 and *Rep* were between 0.520 and 0.821. It was observed that model with LP1 and LP2 led to higher estimated heritability and lower repeatability than the models with higher order.

**Table 4.**
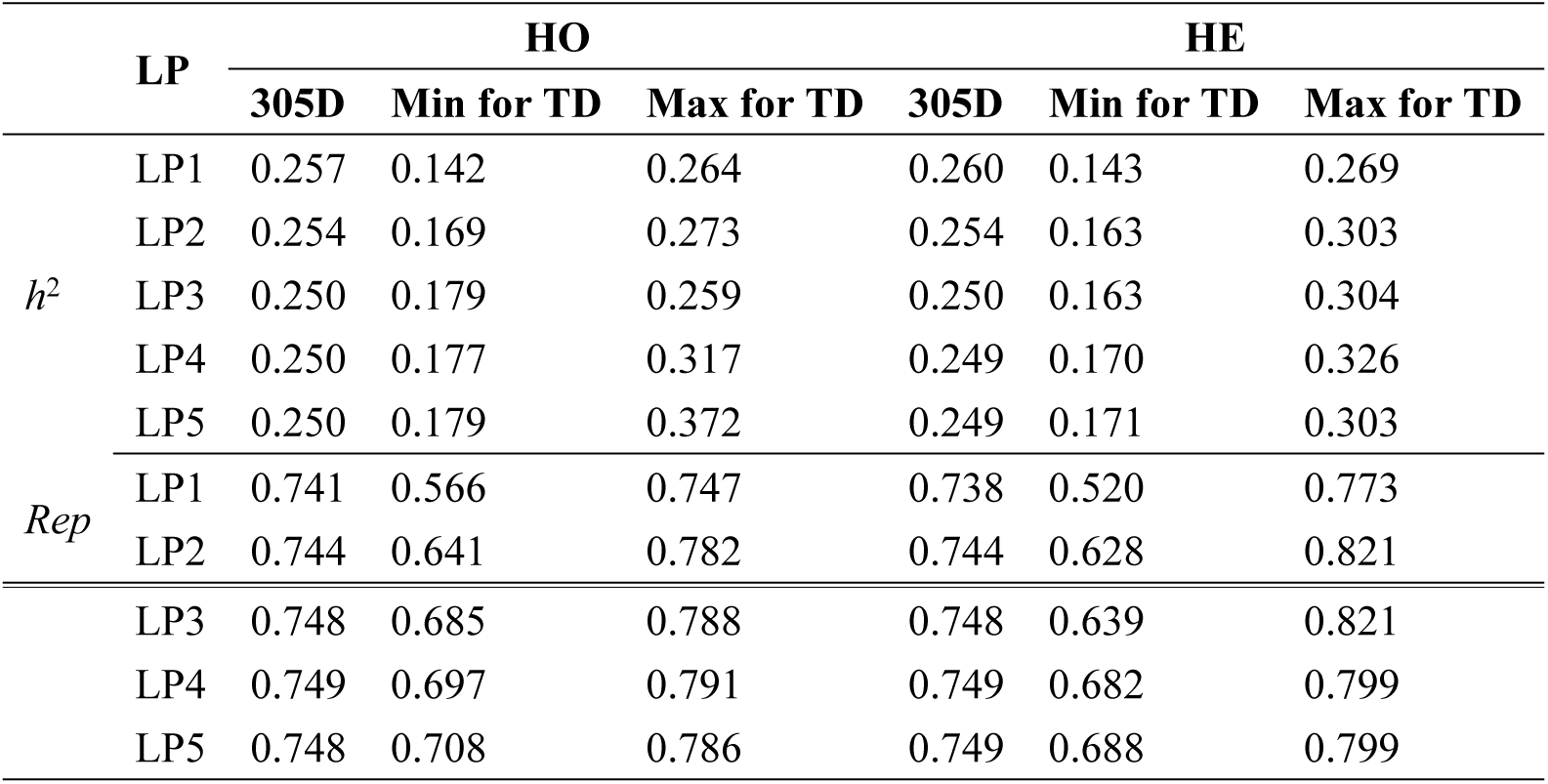
Heritabilities (*h*^2^) and Repeatabilities (*Rep*) for 305d milk yield, and minimal (Min) and maximal (Max) *h*^2^ and *Rep* for TD in different Legendre Polynomials (LP) considering homogeneous (HO) and heterogeneous (HE) residual variances

### Comparison on predictability of models

Spearman’s rank correlations between EBVs from full data and reduced data are shown in Table 5. Correlations for models with different orders ranged from 0.703 to 0.731 and from 0.694 to 0.733 based on homogeneous and heterogeneous residual variances, respectively. Correlations between EBVs increased from LP1 to LP3 and then decreased from LP3 to LP5. The changes in correlation were the same for models with homogeneous and heterogeneous residual variances. The highest correlation of EBVs was found from LP3 and then followed by LP4 and LP5, the lowest correlations between EBVs was from LP1.

**Table 5.** Spearman’s rank correlations between *EBVs* for 305d obtained from full and reduced data for models with different order of Legendre Polynomials (LP) and with homogeneous (HO) or Heterogeneous (HE) residual variance

| **Residual**<br>**Variance** | **LP1** | **LP2** | **LP3** | **LP4** | **LP5** |
| --- | --- | --- | --- | --- | --- |
| **HO** | 0.703 | 0.713 | 0.731 | 0.730 | 0.729 |
| **HE** | 0.694 | 0.711 | 0.733 | 0.732 | 0.729 |

## DISCUSSION

In this study, various criteria were used to compare random regression test day models with different order of Legendre polynomials. Comparison criteria for models have been discussed by (20–22). In general, the smaller 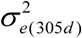 and *-2LogL*, the better goodness of fit for models. In our study, models with higher order obtained lower 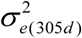 and *-2LogL* values, which was in line with previous studies (17). This means model with LP5 fits data best in terms of residual variance and the likelihood-ratio test. However, models with higher orders introduced more parameters resulting in higher computational demanding (20). Therefore, model selection needs to balance between goodness of fit and computational requirement.

Furthermore, the improvements of goodness of fit became smaller as increasing order of LP in model. For example, the reduction in 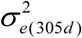 was 14.50% (13.87%), 7.96% (8.90%), 4.67% (3.72%), and 4.06% (4.92%) when comparing successive order LP2, LP3, LP4, LP5 for homogeneous (heterogeneous) residual variances respectively, and for *-2LogL*, the reduction was 1.05% (0.02%), 0.40% (0.03%), 0.19% (0.03%) and 0.12% (0.03%) correspondingly. Especially, there were smaller improvement from LP3 to LP4 and from LP4 to LP5 based on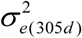 or *-2LogL*. Similar reductions were observed by (23, 24). This means it might be enough to select LP3 or LP4 based on using Comparison criteria and computational requirement.

The trajectory of additive genetic variances and permanent environmental variances showed a quick decreasing in the beginning of lactation and then increased until end of the lactation. This trend is consistent with previous studies in Chinese Holsteins from (1), Brazilian Holsteins by (17, 25). Particularly, (1) used a random regression test-day model with LP4 to estimate parameters for milk yield, they found almost the same curves as the current study for additive genetic variances and permanent environmental variances in Chinese first lactation Holstein cows.

Higher additive genetic variances were in the beginning and end of lactation, which might be attributed to variations in the number of TD records, milk yield level, or non-genetic factors for example pregnancy effects (26). This was coincident with higher genetic variances at the beginning and end of lactation but lower at middle. Other studies have shown that fitting higher order of LP produced higher estimates of genetic variances at the edges of lactation and oscillatory pattern along the lactation trajectory, which might be unlikely biologically (24, 27, 28). This indicates that a model with higher order (e.g., LP5) may not be optimal than a model with lower order (e.g., LP3 or LP4).

The rank correlation of EBVs for 305d from full data and reduced data increased from LP1 (0.703) to LP3 (0.731), then decreased from LP3 to LP5 (0.729). Rank correlations between EBVs using random regression model and EBVs predicted from linear model were between 0.86 to 0.96 for bulls and 0.80 to 0.87 for cows. Random regression model fitted by fourth order Legendre polynomials is recommended for genetic evaluations of Brazilian Holstein cattle (28). In this study, the model with third order and heterogeneous variances had the best predictive ability. Future research should consider using records from different parities and multiple traits such as fat and protein yields.

### Conclusion

This study showed that a random regression test-day model using LP3 or LP4 and accounting for heterogeneous residual variance could achieve reasonable good estimates of variance components. Moreover, model with LP3 and heterogeneous variances had the best prediction ability. This model could be used as an initial model for the implementation of a genetic improvement program in the Chinese Holstein population.

## ACKNOWLEDGMENTS

We acknowledge the funding from Natural Science Foundation of Shandong Province (ZR2016CM37), Key Research and Development Plan of Shandong Province (2018GNC113003), China Agriculture Research System (CARS-36), Agricultural scientific and technological innovation project of Shandong Academy of Agricultural Sciences (CXGC2016A04), and Agricultural improved varieties project of Shandong Province (2016LZGC027), Jinan International Cooperation Plan on Science and Technology (201401353). We thank Dr. Lingzhao Fang and Binjie Li for helping with the calculation of EBV.

